# DESIGN OF A CHIMERIC ACE-2/Fc-SILENT FUSION PROTEIN WITH ULTRAHIGH AFFINITY AND NEUTRALIZING CAPACITY FOR SARS-CoV-2 VARIANTS

**DOI:** 10.1101/2022.06.23.497326

**Authors:** Neil M. Bodie, David Connolly, Jennifer Chu, Bruce D Uhal

**Affiliations:** Paradigm Immunotherapeutics, Monrovia, CA, USA; College of Osteopathic Medicine, Michigan State University, East Lansing, MI, USA; Innovation Lab, ACROBiosystems,1 Innovation Way, Newark, DE, 19711, USA; Department of Physiology, Michigan State University, East Lansing, MI, USA

**Author notes:** Corresponding Author Department of Physiology Michigan State University, 3197 Biomedical and Physical Sciences Building 567 Wilson Road, East Lansing, MI, 48824, USA.

## Abstract

As the coronavirus SARS-CoV-2 continues to mutate into Variants of Concern (VOC), there is a growing and urgent need to develop effective antivirals to combat the newly emerged infectious disease COVID-19. Recent data indicate that monoclonal antibodies developed early in the pandemic are no longer capable of effectively neutralizing currently active VOCs. This report describes the design of a class of variant-agnostic chimeric molecules consisting of an Angiotensin Converting Enzyme-2 (ACE-2) domain mutated to retain ultrahigh affinity binding to a wide variety of SARS-CoV-2 variants, coupled to an Fc-silent immunoglobulin domain that eliminates antibody-dependent enhancement (ADE) and simultaneously extends biological half-life compared to existing mABs. Molecular modeling revealed that ACE-2 mutations L27, V34 and E90 resulted in ultrahigh affinity binding of the LVE-ACE-2 domain to the widest variety of VOCs, with KDs of 93 pM, 507 pM and 73 pM for binding to the Alpha B1.1.7, Delta B.1.617.2 and Omicron B.1.1.529 variants, and notably, 78fM affinity to the Omicron BA.2 variant, respectively. Surrogate viral neutralization assays (sVNT) revealed titers of <u>≥4.9ng/ml</u>, for neutralization of recombinant viral proteins corresponding to the Alpha, Delta and Omicron variants. The values above were obtained with LVE-ACE-2/mAB chimeras containing the Y-T-E sequence that enhances binding to the FcRn receptor, which in turn is expected to extend biological half-life 3-4-fold. It is proposed that this new class of chimeric ACE-2/mABs will constitute variant-agnostic and cost-effective prophylactics against SARS-CoV-2, particularly when administered by nasal delivery systems.

## 1.1 Introduction

SARS-CoV-2 has caused the pandemic Coronavirus Disease 2019 (COVID-19), a highly infectious, fatal disease that affects the lungs and other organs. SARS-CoV-2 belongs to the large coronavirus (CoV) family, which are enveloped viruses that have a 26-32 kb, positive-sense, single-stranded RNA genome (Samavati and Uhal, 2020). The viral envelope consists of a lipid bilayer where the viral membrane (M), envelope (E) and spike (S) structural proteins are anchored. The S protein, also known as viral fusion protein, specifically interacts with its primary receptor, the angiotensin-converting enzyme 2 (ACE-2) on the cell surface to mediate virus-cell fusion, resulting in viral infection (Heurich et al, 2014). The viral S-protein binds to its primary receptor ACE-2 through the Receptor Binding Domain (RBD) of the S subunit (Seyran et al., 2020). A subset of the viral mutations that differentiate variants of SARS-CoV-2 occurs in the RBD portion of the S-subunit (Celik et al., 2022) and thereby affect binding affinity to the viral receptor. In general, mutations that increase binding affinity of the RBD to ACE-2 result in higher infectivity, but other factors such as immune evasion also play important roles in SARS-CoV-2 virulence (Oh et al., 2022).

There is an urgent need to develop effective antivirals to combat this newly emerged infectious disease. At the time of this writing, four monoclonal antibodies have been granted Emergency Use Authorization (EUA) by the US FDA for clinical use: 1) bamlanivimab plus etesevimab are neutralizing mAbs that bind to different, but overlapping, epitopes in the spike protein RBD; 2) Casirivimab plus imdevimab (REGEN-COV) are recombinant human mAbs that bind to nonoverlapping epitopes of the spike protein RBD; 3) tixagevimab plus cilgavimab (Evusheld) are recombinant human anti-SARS-CoV-2 mAbs that bind nonoverlapping epitopes of the spike protein RBD, and 4) sotrovimab targets an epitope in the RBD that is conserved between SARS-CoV and SARS-CoV-2.

Although initially effective at viral neutralization, mAB combinations 1) and 2) have been shown to have significantly reduced neutralizing capacity for the Omicron variant (Shah et al., 2022; Tatham et al., 2022); indeed, on January 24, 2022 the FDA revoked the Emergency Use Authorization (EUA) for REGEN-COV against the Omicron Variant Of Concern (VOC, REGEN-COV Usage Revisions, regencov.com) due to a lack of efficacy against Omicron, and the efficacy of mAB combination 3) for the Omicron variant, initially deemed unclear (Planas et al., 2021), was recently (February 23, 2022) found by the FDA (<u>About EVUSHELD</u>)to be reduced to the point of requiring a doubling of the dose of Evusheld used to combat the Omicron VOC. Although sotrovimab was thought to retain significant capacity to neutralize the initial Omicron variants at the time of this writing (Cameroni et al., 2021), the FDA subsequently revoked the EUA for sotrovimab on April 5, 2022 (https://www.sotrovimab.com), due to loss of efficacy against Omicron BA.2. Therefore, new variant-agnostic approaches to viral neutralization are needed for the current and future pandemics.

Several research groups have developed mutated ACE-2 mimics, including engineered ACE-2 with optimized binding to the viral RBD (Chan et al., 2020, Glascow et al., 2020) and an ACE-2 triple decoy with enhanced affinity for viral variants (Tanaka et al., 2021). Although each of the above engineered ACE-2 mimics had higher affinity binding to the SARS-2 RBD than wild-type ACE-2, they all have nanomolar or low, sub-nanomolar binding affinities. In contrast, the data reported herein show binding of our optimized ACE-Fc construct in the picomolar and even femtomolar range to the Omicron BA.1 and BA.2 variants, respectively. In addition, none of the above constructs incorporate the Fc-silent technologies to be discussed below.

In addition to resistance of existing products to newly emerging variants of SARS-CoV-2, the concept of Antibody-Dependent Enhancement (ADE) of viral infection is increasingly being recognized as a serious danger in COVID-19 (Sánchez-Zuno et al., 2021, Maemura et al, 2021). In ADE, which was observed in previous outbreaks of dengue virus (DENV), Zika virus (ZIKV), Ebola virus, human immunodeficiency virus (HIV), Aleutian mink disease parvovirus, Coxsackie B virus and others, pre-existing non-neutralizing or sub-neutralizing antibodies against viral surface proteins, generated during a previous infection, promote the subsequent entry of viruses into the cell during a secondary infection, and thereby intensify the ensuing inflammatory process (12).

However, ADE has not yet been clinically demonstrated in SARS CoV-2 infection. Instead, “Antibody Dependent Inflammation” (ADI) has been documented (Junqueire et al., 2022). Although severe COVID-19 disease is linked to exuberant inflammation, how SARS-CoV-2 triggers inflammation is not well understood. Monocytes and macrophages are sentinel cells that sense invasive infection to form inflammasomes that activate caspase-1 and gasdermin D (GSDMD), leading to inflammatory death (pyroptosis) and release of potent inflammatory mediators. Junqueire et. al. showed that about 6% of blood monocytes in COVID-19 patients are infected with SARS-CoV-2. Monocyte infection depends on uptake of antibody-opsonized virus by Fcγ receptors, but vaccine recipient plasma does not promote antibody-dependent monocyte infection. SARS-CoV-2 begins to replicate in monocytes, but infection is aborted, and infectious virus is not detected in infected monocyte culture supernatants. Instead, infected cells undergo inflammatory cell death (pyroptosis) mediated by activation of NLRP3 and AIM2 inflammasomes, caspase-1 and GSDMD. Moreover, tissue-resident macrophages, but not infected epithelial and endothelial cells from COVID-19 lung autopsies, have activated inflammasomes. These findings, taken together, suggest that antibody-mediated SARS-CoV-2 uptake by monocytes/macrophages triggers inflammatory cell death that aborts production of infectious virus, but causes systemic Fc receptor-dependent inflammation that contributes to COVID-19 pathogenesis (Junqueire et. al., 2022).

For all the reasons above, the objective of this work was to design molecules with high but variant-agnostic binding affinities to a wide variety of SARS-CoV-2 variants, particularly if they might also incorporate features that reduce or eliminate ADI or ADE. In this report we describe the design of just such molecules, which combine a synthetic human ACE-2 domain containing mutations that allow variant-agnostic, ultra-high affinity binding to the SARS-CoV-2 S1 subunit, combined together with an Fc-silent antibody domain that essentially eliminates the potential for ADI or ADE. Moreover, a third mutant option of the antibody domain of the chimera is offered, with the intent to substantially increase (3-4-fold) the biological half-life of the chimera if delivered by aerosol or nasal administration. Given that nasal infection is the most likely route of SARS-CoV-2 entry in humans (Hou et al., 2020), it is proposed that a nasal administration of the new chimeric molecules described herein will constitute an effective prophylactic against SARS-CoV-2 infection that will not only be effective, but also is expected to be economically far superior to current monoclonal antibody treatments for COVID-19.

### 1.2 Materials and Methods

#### 1.2.1 Viral and protein constructs

Viral Receptor Binding Domain (RBD), S1 protein subunit and whole spike protein trimers were synthesized by ACRO Biosystems (Newark, DE) as recombinant proteins designed on the basis of publicly available sequence data (outbreak.info). Recombinant human ACE-2 constructs were synthesized by Absolute Antibody (Boston, MA) designed on the basis of sequence data obtained from Uniprot, ACE2 (Gln18-Ser740) Accession #Q9BYF1 protein sequence data and modified as described in Figures 2-5. GenScript ACE2 (Gln18-Ser740) Accession #Q9BYF1 Fc Chimera (Cat. No. Z03516) was purchased from GenScript (Piscataway NJ). The ACE-2/mAB chimeras were synthesized by Absolute Antibody (Wilton Centre, UK).

**Figure 1.**
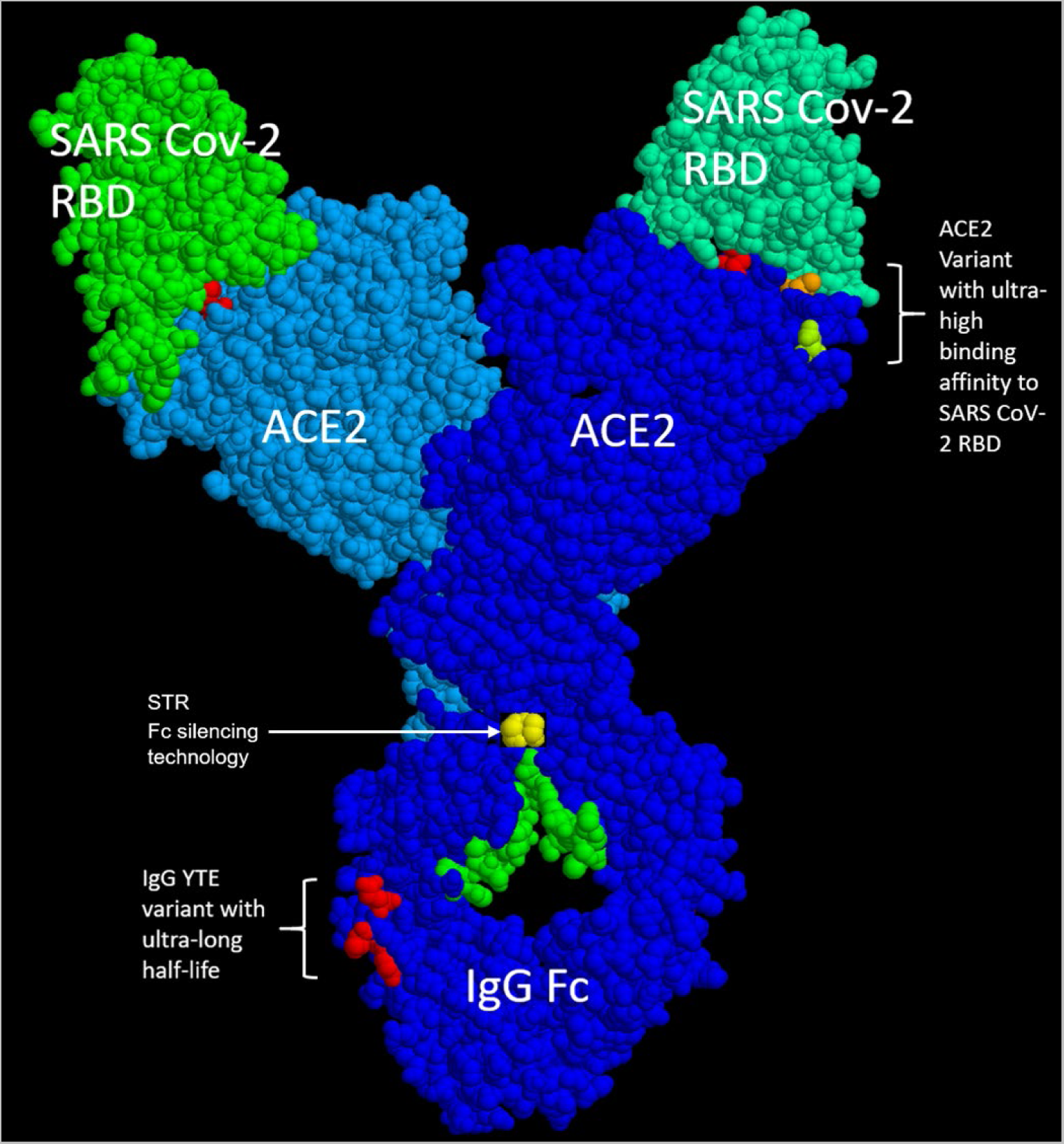
Critical features of Paradigm’s chimeric ACE-2/Fc-silent antibody technology. A: The use of a full length (amino acids 18-740) ACE-2 (blue), which substantially increases binding affinity for SARS CoV-2 RBD/spike protein (green). B: the construct of a mutated ACE-2 variant (top right red, orange, light green amino acids) with picomolar affinity for the RBD and ∼femtomolar affinity for the full-length spike protein. C: linkage of the ACE-2 construct to a completely Fc-silent “STR” mutated monoclonal antibody (yellow, the model was generated with RASMOL using an IgG Fc, Mike Clark Ph.D., used with permission); the use of STR technology in an ACE2 chimeric prophylactic for SARS CoV-2 is patent pending, see Fig.12. D: one of the two mAb chimeras (“LiVE-Longer” vs. “LiVE”) utilizes a Y-T-E variant (red amino acids, bottom) for increased half-life by binding to FcRn, which recycles IgG and is thereby predicted to increase its biological half-life by 3-4-fold (see Fig.13 and text for details). The “LiVE” mAB does not contain the YTE sequence; both LiVE and LiVE-Longer are tested in Fig.11 below and Table I. This figure was generated by Protean 3D, Version 17.3 (DNASTAR. Madison, WI).

**Figure 2.**
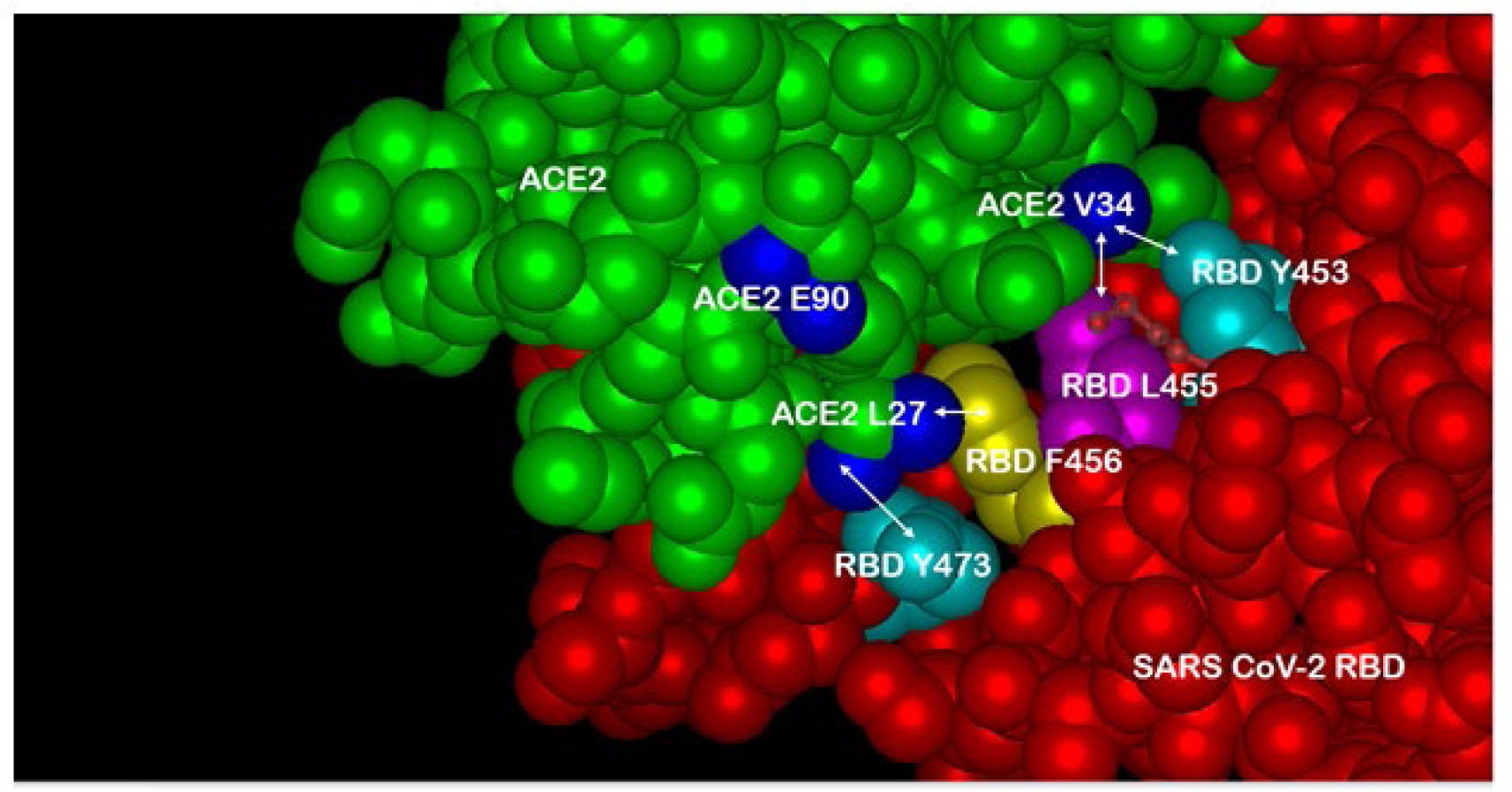
Three-dimensional model of the SARS-CoV-2/ACE-2 molecular interface. Multiple In Silico modeling programs were used to predict optimal amino acid substitutions in ACE-2 (green) that would retain high binding affinity to the widest possible range of amino acid variations in the SARS-CoV-2 spike protein receptor binding domain (RBD, red). Two of the three optimal substitutions in ACE-2 depicted here are: T27L (ACE2 L27, blue lower) and H34V (ACE2 V34, blue upper right), shown here in close proximity to RBD amino acids Y473 and F456 versus L455 and Y453, respectively. The third substitution N90E (ACE2 E90, blue upper left) is discussed in Figure 3. See text for details.

#### 1.2.2 In silico molecular modeling

Molecular models were derived from publicly available ACE-2 and SARS-CoV-2 sequence databases (Outbreak.info) with Protean 3D Version 17.3 (DNASTAR. Madison, WI) software using the method described by Zhang and Zhang (Zhang and Zhang, 2010), which uses a knowledge-based potential to solve protein folding and protein structure prediction problems. Based on the parent software I-TASSER (Zhou and Skolnick, 2007), the method of Zhang and Zhang can differentiate well between Leucine and Isoleucine, an ability important for the potential analysis of Leucines in viral or ACE-2 variants. Comparisons of atomic-level structures and viral-ACE-2 interactions were achieved with the knowledge-based atomic potential algorithm DFIRE (Zhou and Zhou, 2002), which generates a numerical protein-protein interaction score that becomes more negative with more stabilizing molecular interactions. Mutations in ACE-2 yielding the highest affinity binding to the widest variety of SARS-2 variants were sought. To find these, dozens of in silico experiments were performed to determine the optimal ACE2 mutations (27L and 34V) for the most highly conserved SARSCoV2 RBD anchor amino acid residues RBD L455, F456 and Y473, based on DMS of the SARSCoV2 RBD, see Figure 3B (Starr et al., 2020). The final mutation we chose, ACE2 N90E, was based on the DNASTAR modeling program suggesting steric hinderance of the ACE2 N90 glycan. Aspartic acid at the ACE N90E was chosen over other amino acid residues based on a possible stabilizing interaction with ACE2 K26 predicted by the DNASTAR modeling program.

**Figure 3.**
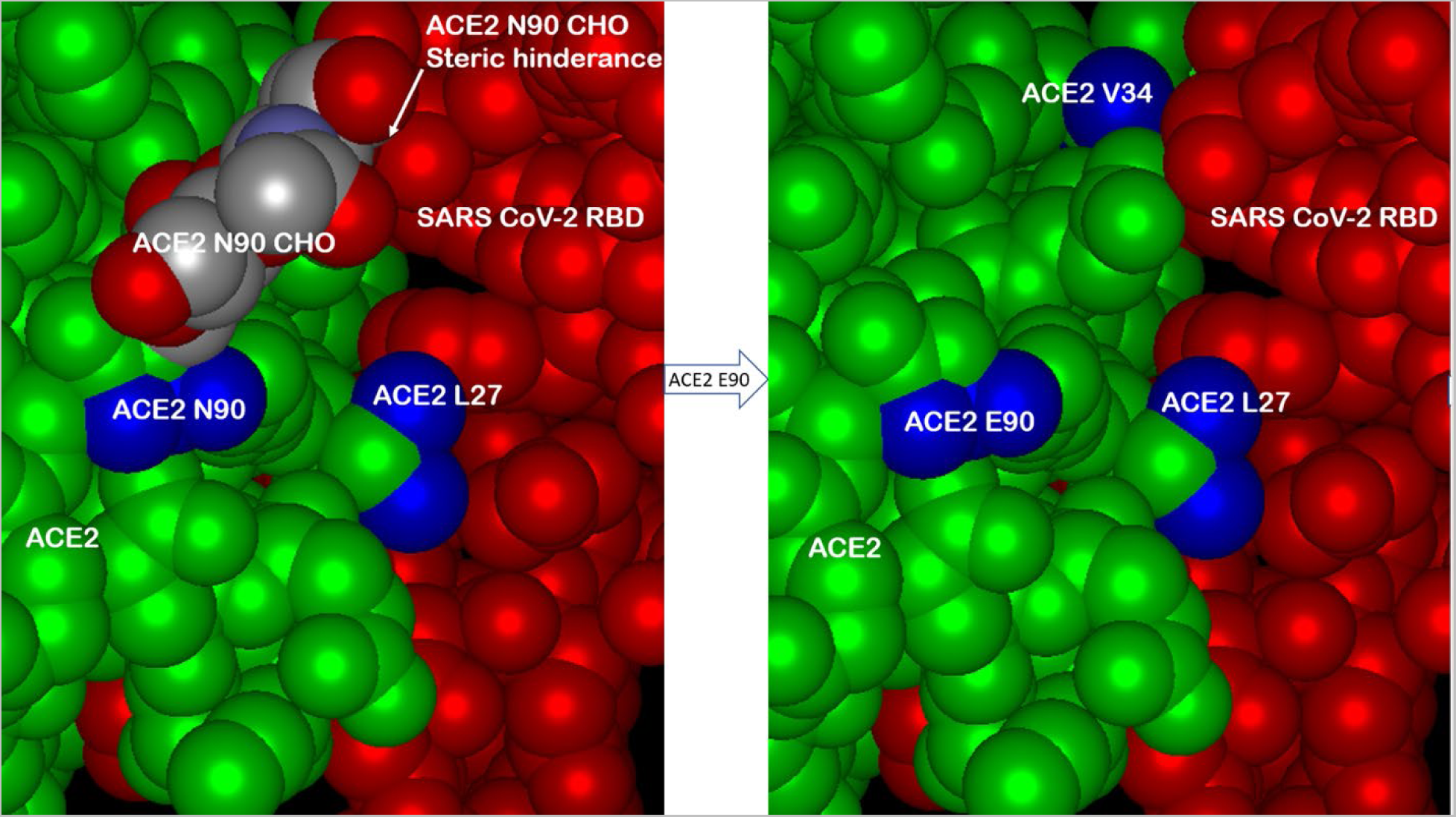
The effect of eliminating the glycosylation site at amino acid 90 of ACE-2. The asparagine at amino acid 90 of wild type ACE-2 (N90) is a site for N-linked glycosylation (left panel, ACE-2 N90 CHO) which, when the sugar is present, causes steric hindrance of ACE-2/RBD interaction. Multiple modeling platforms predicted that elimination of the glycosylation site by the mutation N90E (asparagine to glutamate) would relieve steric hindrance and allow closer ACE-2/RBD interaction, particularly in combination with the ACE-2 H34V substitution (right panel, top). Subsequent figures below show data consistent with this prediction. See text for details.

**Figure 4.**
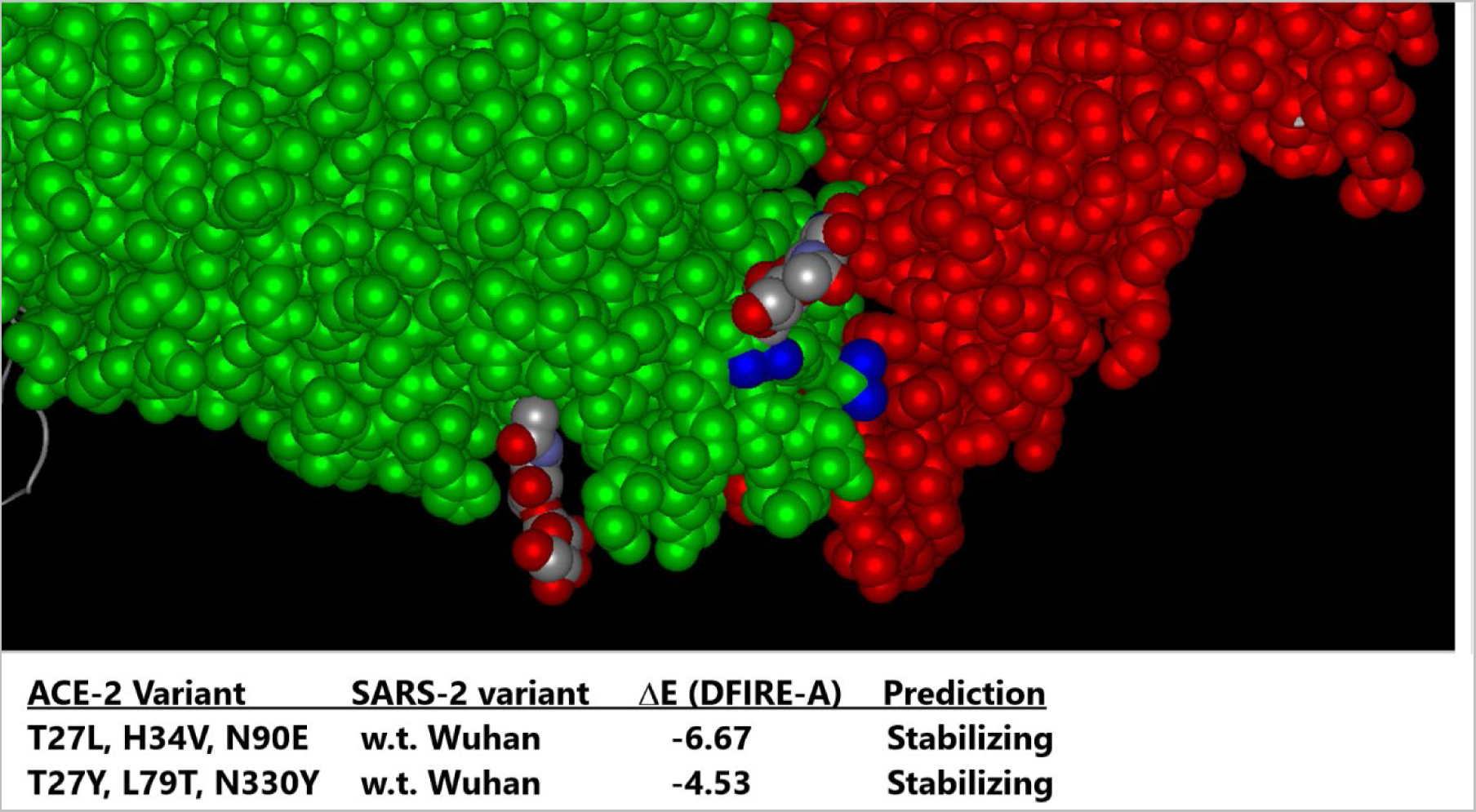
The designed ACE-2 variant LVE/STR chimera scores higher than other ACE-2 Fusion protein competitors by the DFIRE scoring method. Molecular modeling was conducted with Protean 3D Version 17.3 (DNASTAR. Madison, WI) software, aimed at testing Paradigms’ proprietary LiVE variant (ACE-2 mutations T27L, H34V and N90E) binding to w.t. SARS-2 RBD (DFIRE = -6.67), as it compares to the ACE-2 variant YTY (ACE-2 mutations T27Y, L79T and N330Y) binding to the w.t. SARS-2 RBD (DFIRE = -4.53). A lower DFIRE score predicts tighter (stabilizing) protein-protein interactions. See Methods for details.

#### 1.2.3 Binding affinity determinations

Surface Plasmon Resonance (SPR) assays of protein-protein interactions were performed by Acro Biosystems (Newark, DE) on a Biacore T200 Instrument fitted with Series CM5 Sensor Chip. Prior to SPR assay, samples were desalted on Zeba Spin 7K MWCO columns. Binding affinities were determined in HBS-N buffer, 10X (0.1M HEPES, 1.5MNaCl) containing EDTA and Tween 20, at a flow rate of 30 μL/minute, run for 120 seconds association and 180 seconds dissociation. The reference subtracted SPR binding curves were blank subtracted, and curve fitting was performed with a 1:1 model to obtain kinetic parameters using the Biacore T200 Evaluation software. Binding data are reported as estimated dissociation constant (KD, Figures 6,10 and Table I).

**Figure 5.**
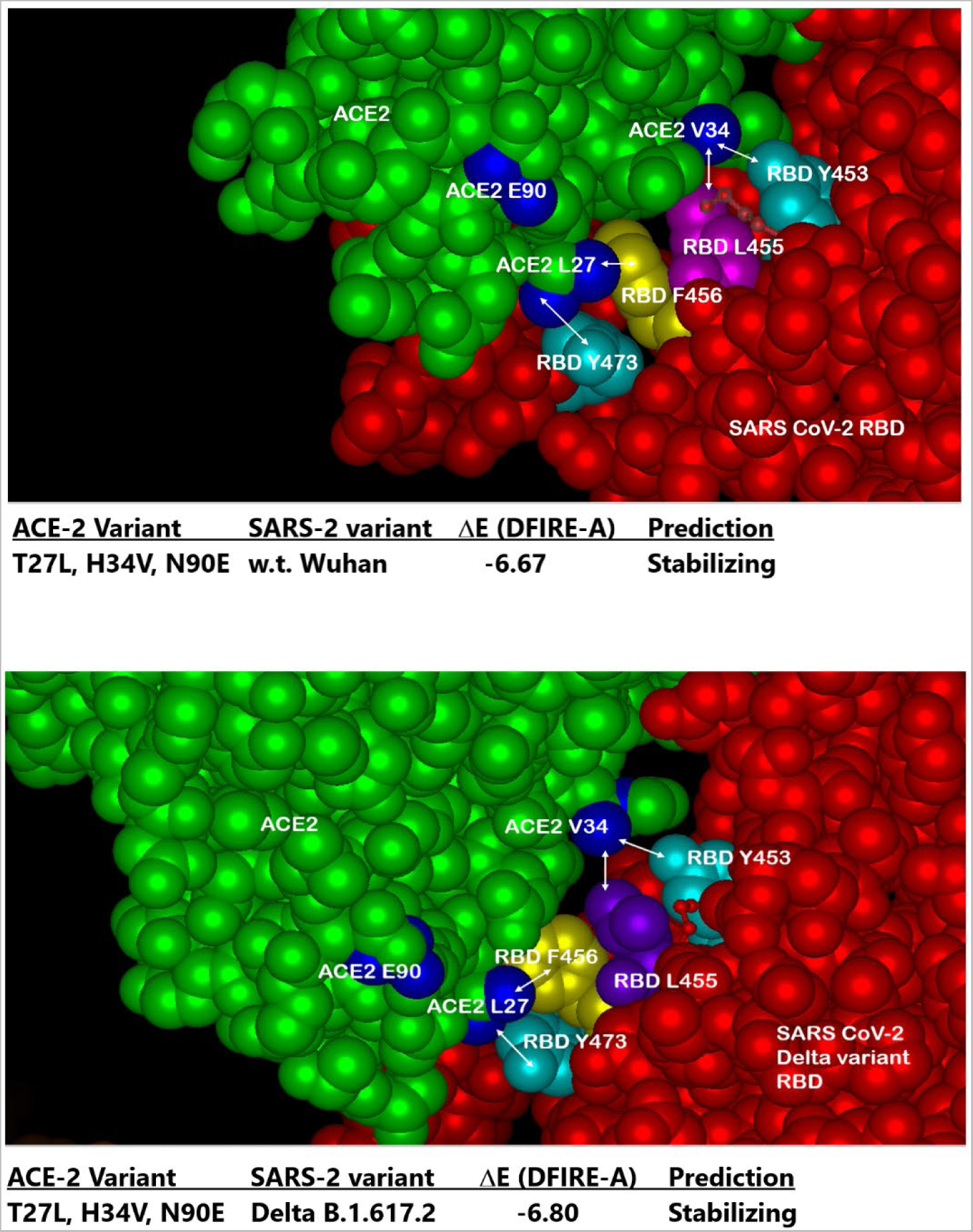
In Silico testing of SARS CoV-2 Delta variant predicts Paradigm’s ACE-2 variant LVE/STR chimera out-scores competing ACE-2 chimeric technologies. Top Panel: Molecular modeling with Protean 3D Version 17.3 (DNASTAR. Madison, WI) software of ACE-2 LVE construct (green) binding to w.t. SARS-CoV-2 RBD (red) yields DFIRE score = -6.67. Bottom Panel: Identical modeling of ACE-2 LiVE construct (green) binding to SARS-CoV-2 Delta variant yields DFIRE score = -6.80, slightly tighter (stabilizing) than that for the w.t. RBD. Note rotation of RBD F456 (yellow) in Delta variant toward ACE-2 amino acid 27, made permissive by the T27L mutation.

**Figure 6.**
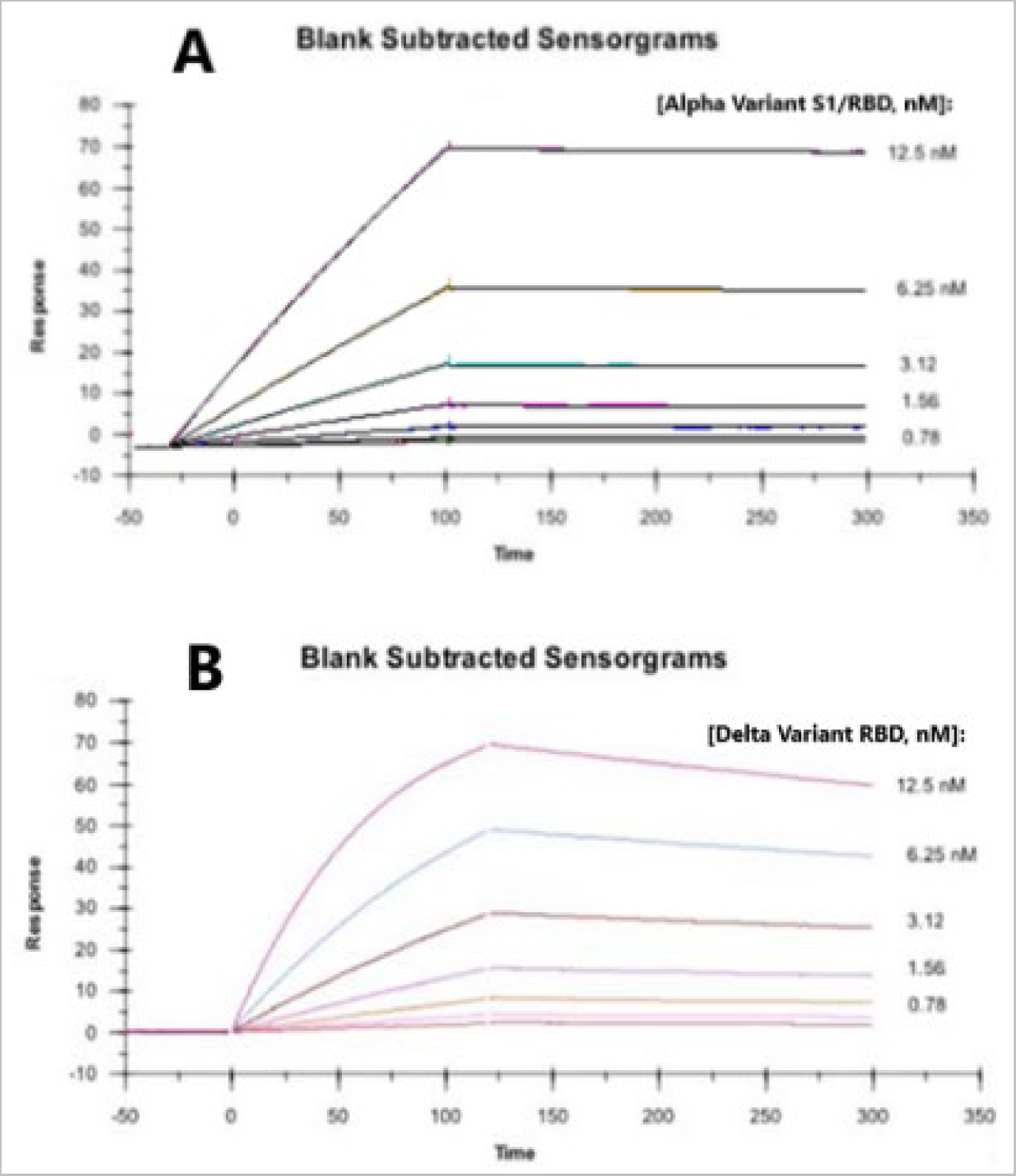
High binding affinity of Paradigm’s ACE-2 variant LVE/STR chimera to the RBD of Alpha and Delta variants of SARS-CoV-2. Surface Plasmon Resonance (SPR) assays were performed to determine binding affinities between the synthesized, purified LVE/STR chimera and synthesized RBDs of SARS-CoV-2 Alpha or Delta variants. Panel A: The S1 subunit with RBD of SARS-CoV-2 Alpha variant B.1.1.7 was synthesized, purified and subjected to SPR assay against the purified LVE/STR chimera. Determined binding affinity was 0.378 nM (Acro Biosystems). Note slow “off rate” compared to Panel B below. Panel B: The RBD of Delta variant B.1.617.2 was synthesized, purified and subjected to SPR assay against the purified LiVE/STR chimera. Determined binding affinity was 0.554 nM (Acro Biosystems). See Table I for more complete listing of binding affinities.

#### 1.2.4 Surrogate viral neutralization tests

The surrogate Viral Neutralization Test (sVNT, C-PASS, GenScript USA Inc., Piscataway, NJ 08854) was used to characterize the antibodies described, using the methodology defined previously (Tan et al., 2020). The manufacturer’s instructions were followed without modification except where specifically noted below. Briefly, the SARS-CoV-2 sVNT Kit is a blocking ELISA detection tool, which mimics the virus neutralization process. The kit contains two key components: the Horseradish peroxidase (HRP) conjugated recombinant SARS-CoV-2 RBD fragment (HRP-RBD) and the human ACE2 receptor protein (hACE2). The protein-protein interaction between HRP-RBD and hACE2 can be blocked by neutralizing antibodies against SARS-CoV-2 RBD; specific applications are described in the Legends to Figures.

### 1.3 Results

The basic design of Paradigm’s ACE-2-Fusion protein chimera is shown in Figure 1. A synthetic, mutant variant of human ACE-2 was designed to have high-affinity binding to a wide variety of SARS-CoV-2 variants including Delta and Omicron (data shown below). The ACE-2 was linked to an Fc-silent designer monoclonal antibody of two forms: Fc-silent (L234S/L235T/G236R, “STR”, Wilkinson et al., 2021) and a “live longer” Fc-silent version containing the YTE variant (M252Y/S254T/T256E, Dall’Acqua et al., 2006), which increases binding to FcRn receptor to achieve longer biological half-life as discussed below. Figure 2 shows a three-dimensional model of the ACE-2/SARS-CoV2 molecular interface with the ACE-2 mutations found to impart high binding affinity to the widest range of SARS-2 variants. In particular, the amino acid (AA) substitutions T27L and H34V interact with SARS-2 RBD amino acids 473 and 456 versus 455 and 453, respectively. The third substitution N90E (ACE2 E90) is discussed next.

Figure 3 shows the effect of the ACE-2 AA substitution N90E, which eliminates the site for N-linked glycosylation of ACE-2 at the ACE-2/SARS-2 interface. Surprisingly, elimination of the N-linked glycan resulted in higher affinity binding to several SARS-2 variants, presumably due to loss of steric hindrance otherwise caused by the glycan. In Figure 4, molecular modeling with Protean 3D software of ACE-2 variant binding to a purified, recombinant wild type (w.t.) SARS-2 receptor binding domain (RBD) revealed tighter binding of the ACE-2 LVE variant (DFIRE score -6.67), than that of the ACE-2 amino acid substitutions YTY at positions 27,79 and 330, respectively (DFIRE score -4.53). See Methods for details.

Figure 5 shows three dimensional molecular models and binding affinity predictions for the interactions between Paradigm’s ACE-2 LVE/STR chimeric Fusion protein and the purified recombinant RBDs of the w.t. SARS-2 variant (top panel) versus the RBD of the Delta B.1.617.2 variant (bottom panel). The lower DFIRE score of -6.80 for binding to the Delta variant predicts a more stable (tighter) ACE-2/SARS-2 interaction. Consistent with this prediction, the data in Figure 6 show Surface Plasmon Resonance (SPR) data indicating very high binding affinity of Paradigm’s ACE-2 LVE/STR chimera to either the purified S1/RBD of the Alpha B.1.1.7 variant (378pM) or the purified RBD of the Delta B.1.617.2 variant (554 pM).

Surrogate viral neutralization tests (sVNT) are also consistent with the above results; Figure 7 displays sVNT data indicating that Paradigm’s ACE-2 LVE/STR chimeric Fusion protein could neutralize the purified recombinant RBD of either the w.t. SARS-2 Wuhan variant (blue bars) or the Delta variant B1.617.2 (red bars) with nearly equal potency (sVNT titer ∼ 4.9ng/ml). In Figure 8, the LVE/STR chimera neutralized the recombinant RBD of the Beta variant B.1.351 with greater potency (red bars, sVNT titer ∼2.4ng/ml) than that for the RBD w.t. Wuhan variant (blue bars, titer ∼4.9ng/ml). In sVNT tests of the SARS-2 Alpha variant (Figure 9), the LVE ACE-2/STR chimera neutralized the purified RBD of the Alpha variant B1.1.7 significantly better (sVNT titer >4.9ng/ml) than the Genscript Fc-IgG/ACE-2 chimera Z03516 (∼6.3ug/ml).

**Figure 7.**
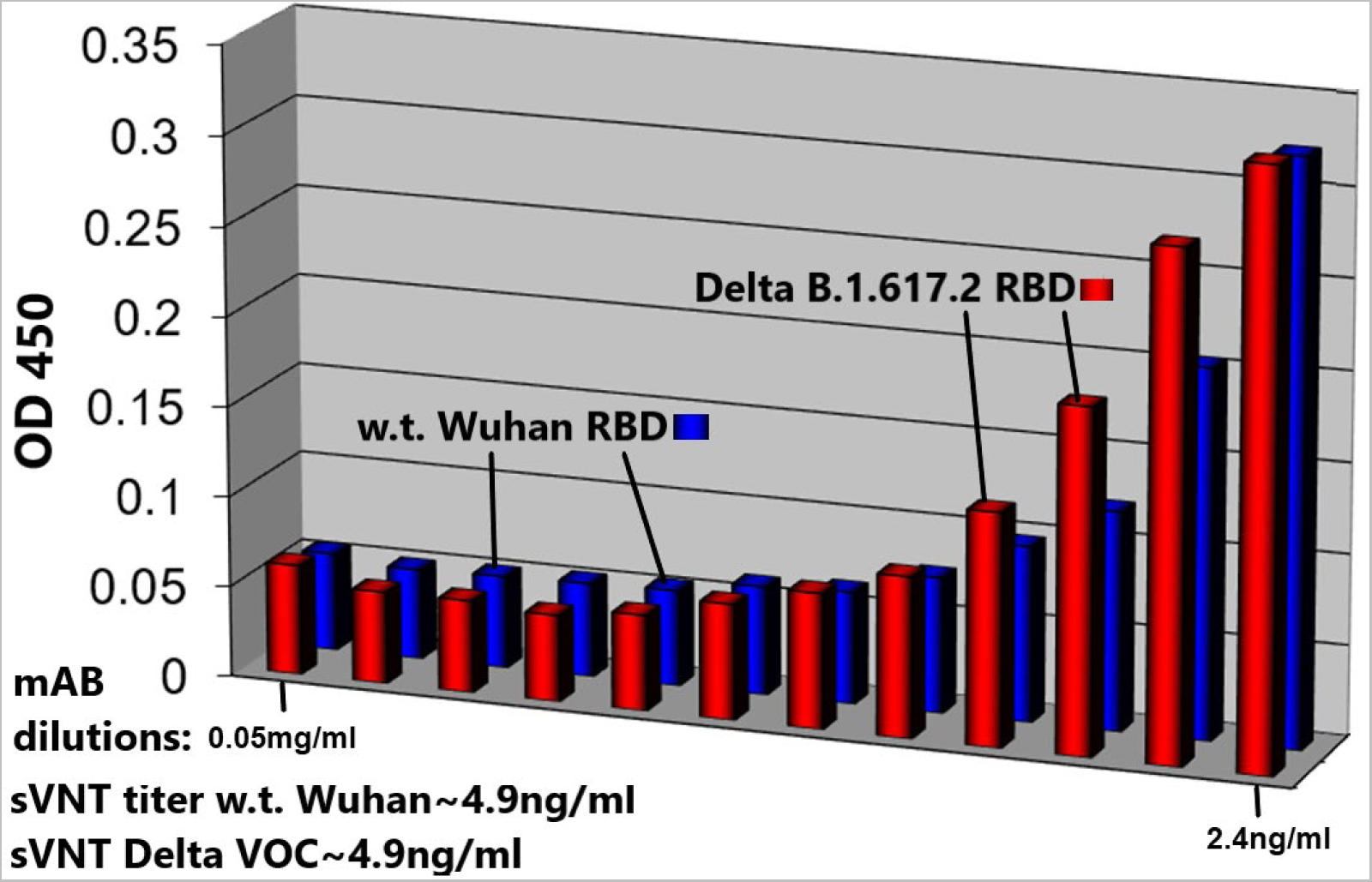
Paradigm’s ACE-2 variant LVE/STR chimera neutralizes Delta Variant B.1.617.2 RBD or w.t. SARS-CoV-2 RBD (Wuhan strain) with nearly equal potency. Surrogate Viral Neutralization Tests (sVNT, see Methods) were performed with SARS-CoV-2 RBD proteins expressing the Delta Variant B.1.617.2 sequence (T478K and L452R mutants, red bars) or the wild type (w.t., original Wuhan strain) sequence (blue bars) incubated with the ACE-2 LiVE/STR construct. Dilutions shown are 0.05mg/ml to 2.4ng/ml, left to right. Note similar sVNT titers of ∼4.9ng/ml.

**Figure 8.**
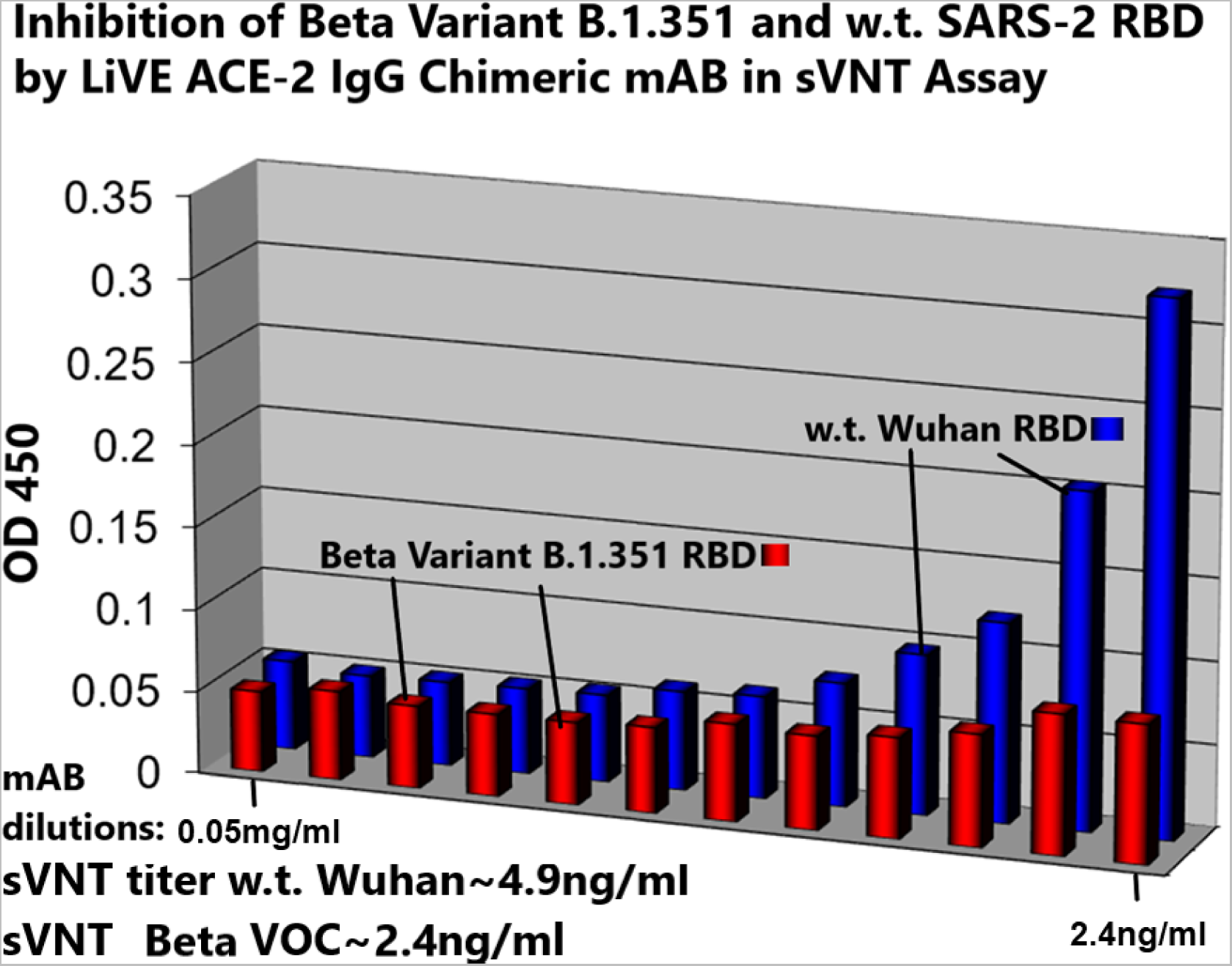
Paradigm’s ACE-2 variant LVE/STR chimera neutralizes Beta Variant B.1.351 RBD significantly better than w.t. SARS-CoV-2 RBD. sVNT assays were performed with SARS-CoV-2 RBD proteins expressing the Beta Variant B.1.351 sequence (K417N, E484K and N501Y mutants, red bars) or the wild type (w.t., original Wuhan strain) sequence (blue bars) incubated with the ACE-2 LiVE/STR construct. Dilutions shown are 0.05mg/ml to 2.4ng/ml, left to right. Note sVNT titers of ∼2.4ng/ml for neutralization of the SARS-2 Beta variant.

**Figure 9.**
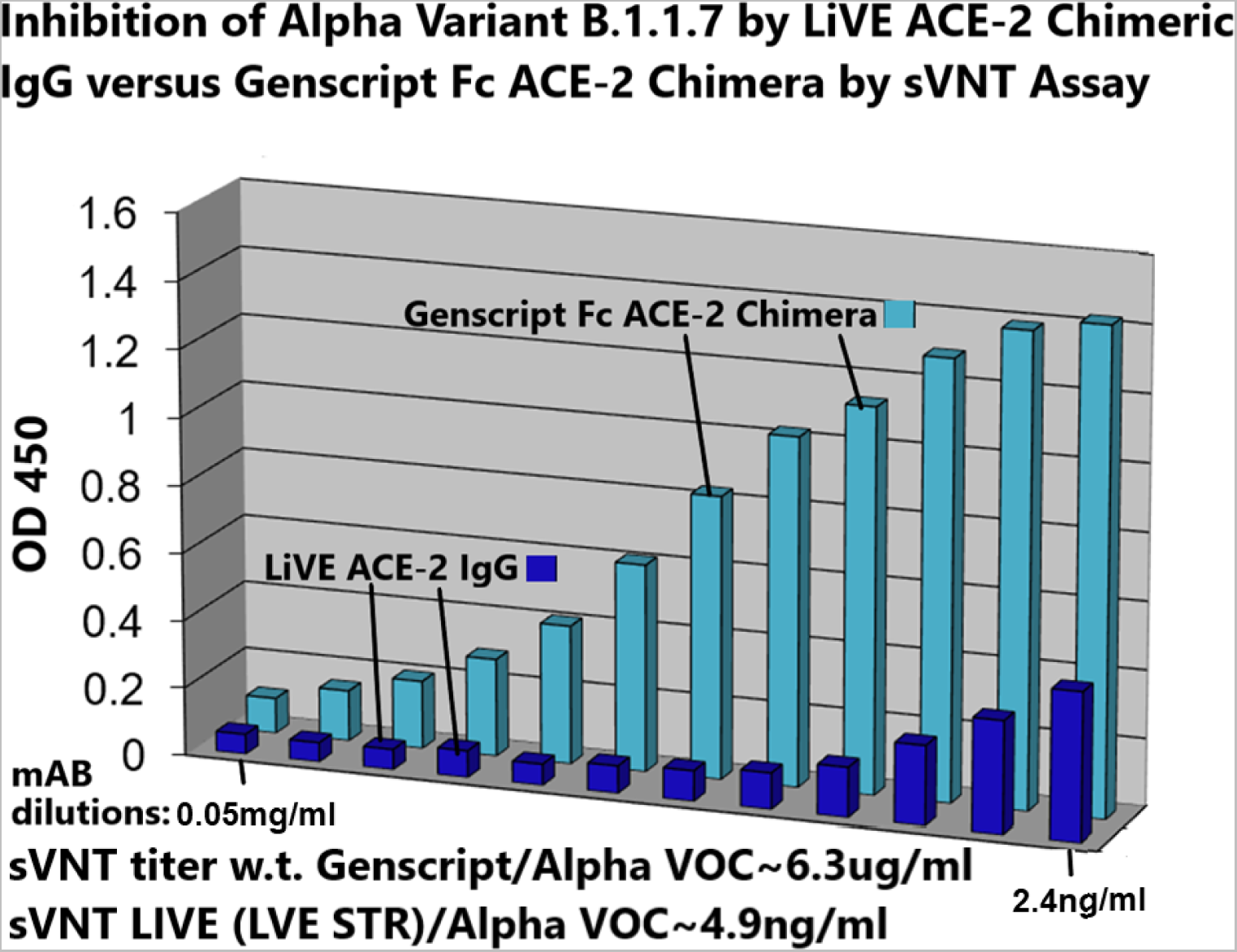
Paradigm’s ACE-2 variant LVE/STR chimera neutralizes Alpha Variant B.1.1.7 RBD significantly better than GenScript IgG FL18-740 w.t. ACE-2 mAB. sVNT assays were performed with SARS-CoV-2 RBD proteins expressing the Alpha Variant B.1.1.7 sequence (N501Y mutant) challenged with either Paradigm’s ACE-2 variant LVE/STR chimera (dark blue bars) or with the GenScript IgG FL18-740 w.t. ACE-2 mAB (light blue bars, binding affinity 3nM). Dilutions shown are 0.05mg/ml to 2.4ng/ml, left to right. Note sVNT titers of ∼6.3ug/ml for the Genscript mAB for neutralization of the Alpha variant, in contrast to ∼4.9ng/ml for neutralization of the Alpha variant by the Paradigm construct LiVE-STR.

**Figure 10.**
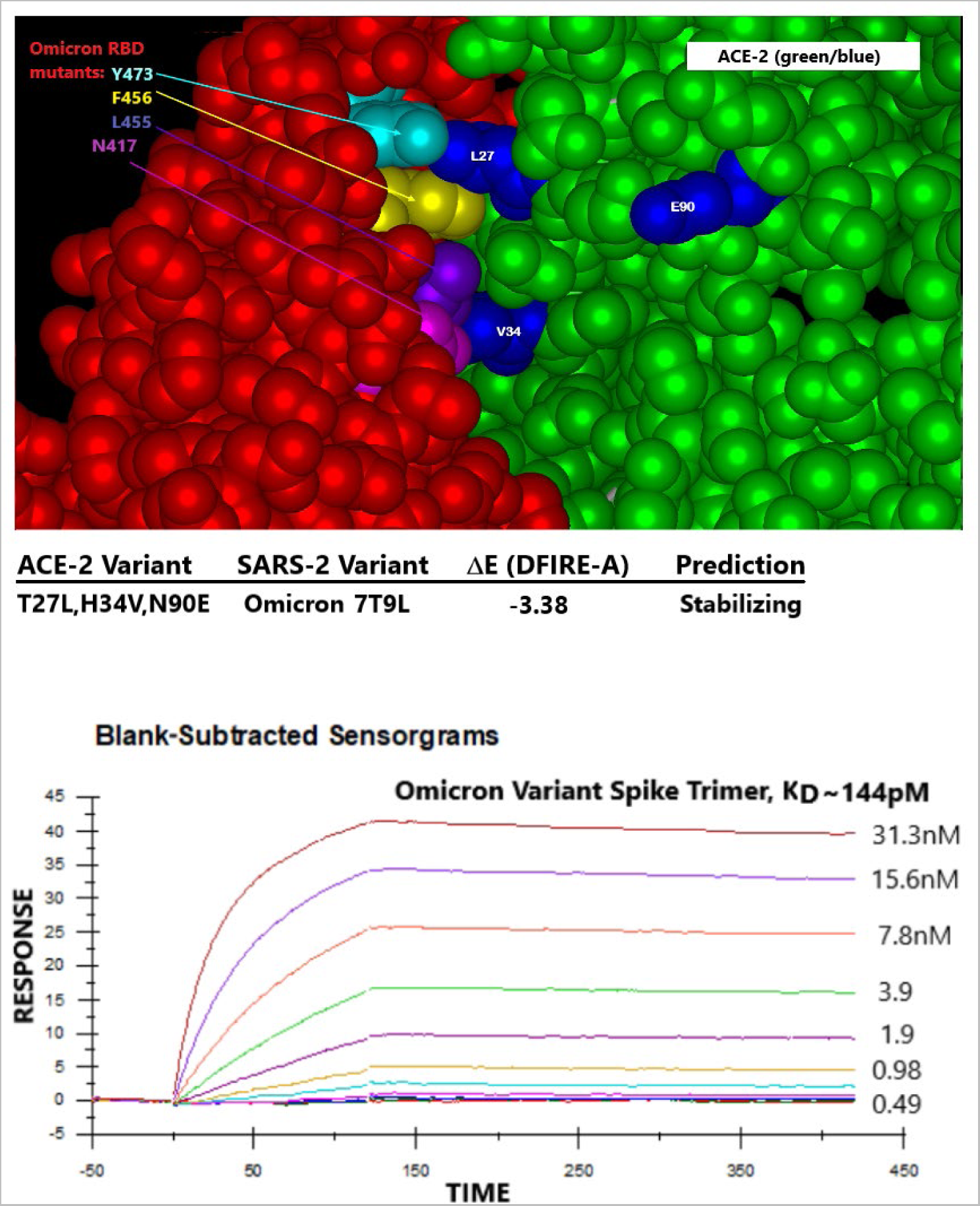
Paradigms’ ACE-2 variant LVE/STR chimera binds to SARS-2 Omicron variant B.1.1.529 spike protein trimer with high affinity. Top Panel: Molecular modeling of Paradigms’ ACE-2 variant LVE/STR chimera binding to the Omicron variant sequence as of 1/11/2022. Omicron RBD (red) mutations Y473, F456, L455 and N417 are shown at upper left, and ACE-2 (green) mutations LVE are shown in blue. The ACE-2 H34V makes contact with the aliphatic straight chain of Omicron mutation Q493R and K417N. Note the vertical orientation of N417 in close contact with V34, which enabled higher affinity binding compared to the w.t. K417, which assumed a more horizontal orientation in earlier SARS-2 variants. Molecular modeling performed identically to that in Figures 4 and 5 yields superior DFIRE score of -3.38 (stabilizing). See Discussion for details. Bottom Panel: The S1 subunit trimer of SARS-CoV-Omicron variant described above was synthesized, purified and subjected to SPR assay against the purified LVE/STR chimera. Determined binding affinity was 0.144nM (Acro Biosystems); see Table I for more complete listing of binding affinities.

Figure 10 displays molecular modeling of the molecular interface between Paradigms’ ACE-2 variant LVE/STR chimera and the RBD of the Omicron variant of SARS-CoV-2 (RBD sequence as of December 12, 2021 at 75% cutoff). Of note, the aliphatic side chain of the Omicron Q493R mutation (purple) makes contact with ACE-2 mutation V34. In molecular modeling (see Methods), simulation of the ACE-2 LVE/STR chimera binding to the Omicron variant yielded a very favorable DFIRE score of -7.26, indicating a tight, stabilizing interaction. This modeling is consistent with the sVNT data in Figure 11, which show that the Omicron variant sequence known on December 12, 2021, when expressed as either a purified recombinant RBD (top panel) or purified spike protein trimer (bottom panel), could be potently neutralized by Paradigm’s “LiVE” ACE-2 variant LVE/STR Fusion protein chimera or by the “LiVE Longer” LVE/STR - YTE Fusion protein chimera (sVNT titers∼4.9ng/ml).

**Figure 11.**
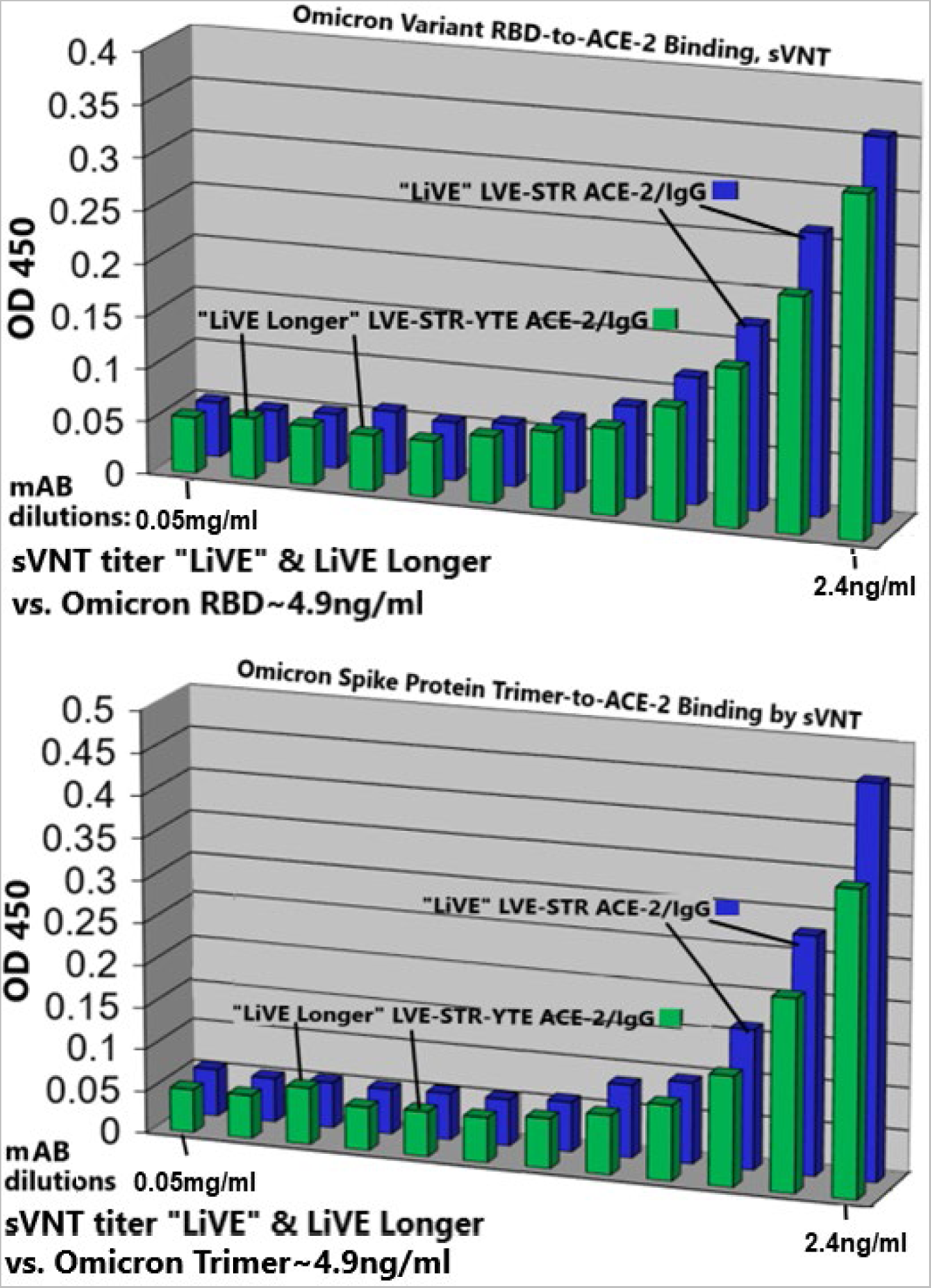
Paradigm’s ACE-2 variant LVE/STR chimeras potently neutralize the SARS-CoV-2 Omicron Variant B.1.1.529. sVNT assays were performed with purified recombinant SARS-CoV-2 RBD (top panel) or spike protein trimers (bottom panel) expressing the Omicron variant sequences described in Fig.10. The bargraphs show inhibition of the binding of Omicron RBD (top) or Omicron spike protein trimer (bottom) to purified recombinant ACE-2 by either Paradigm’s “LiVE” ACE-2 variant LVE/STR STR Fusion protein chimera or by the “LiVE Longer” LVE/STR STR - YTE IgG chimera. Dilutions shown are 0.05mg/ml to 2.4ng/ml, left to right. Note similar sVNT titers of ∼4.9ng/ml for neutralization of Omicron RBD or Omicron spike trimers, and slightly better neutralization by the “LiVE Longer” chimera (green). See Figures 1 and 12 for additional details about the ACE-2/Fusion protein chimeras.

Figure 12 shows the relative potency of Fc-silencing technologies, determined by Surface Plasmon Resonance (SPR) measurements of the binding of purified, recombinant protein samples of each modified antibody to immobilized recombinant human FcγRI receptor. At the far right, both Paradigm’s ACE2 Fc fusion proteins using the STR silencing mutations and mAbsolve’s STR silencing methods show the lowest, almost undetectable binding to FcγRI receptor. The Inset displays SPR data for binding of either the LiVE (non-YTE) or LiVE-Longer chimeras to purified FcRn receptor; note high affinity binding of the YTE chimera to FcRn, but not by the non-YTE variant; binding to FcRn will increase the biological half-life of the chimera in nasal epithelium (see below and Discussion). Figure 13 shows data underlying the inclusion of the YTE mutations in the “LiVE-longer” version of the ACE-2 LVE/STR YTE chimera. Reprinted here with publisher’s permission, Ladel et al. (Ladel et al., 2018) showed that IgG binding to the FcRn receptor, which is enhanced by the YTE mutation, results in slow (4-8 hours) transcytosis and recycling of the IgG in the nasal mucosa, which results in a much longer biological half-life of IgGs administered nasally.

**Figure 12.**
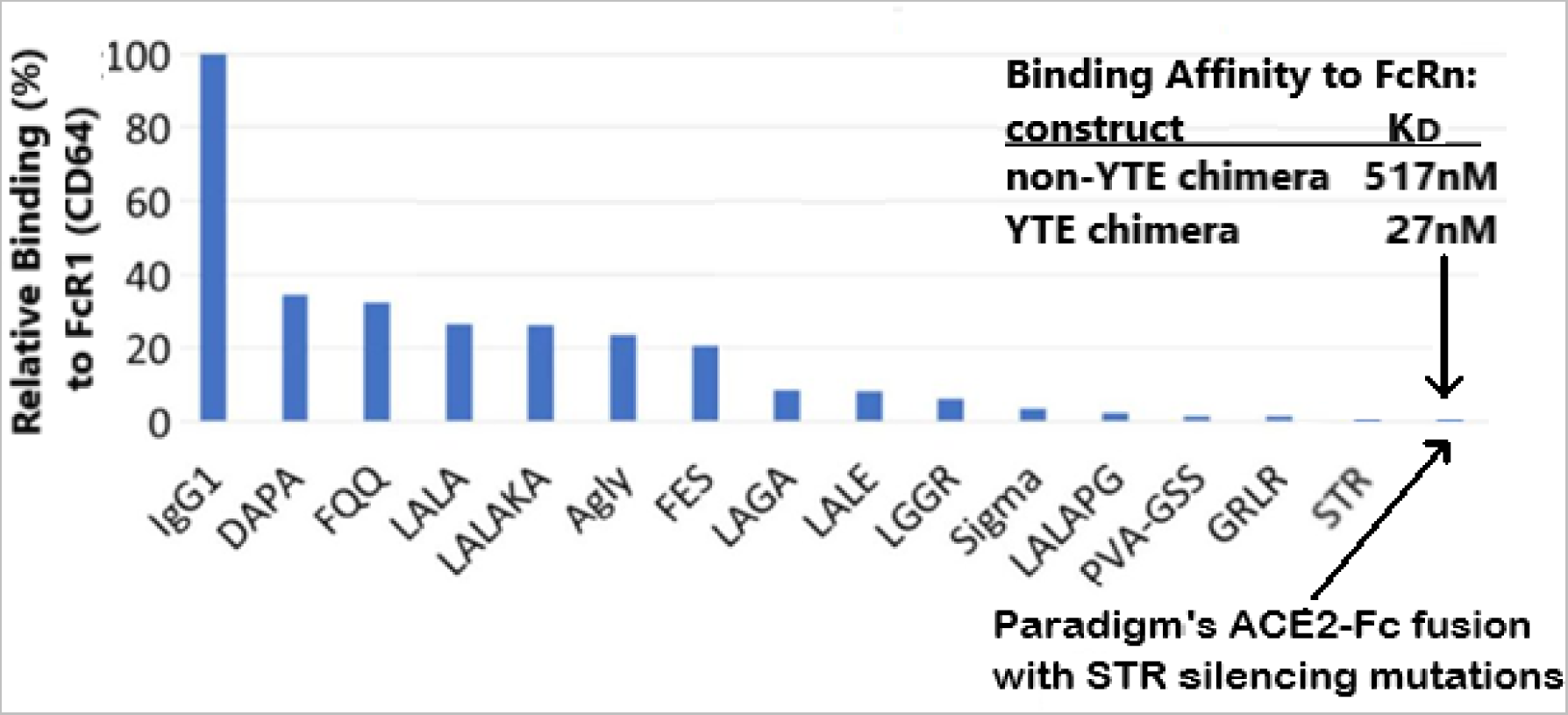
Paradigm’s Fc fusion protein with the STR Fc silencing mutations is Superior to LALA and other Reduced Fc Effector Function Technologies. A comparison of Fc-silencing technologies shows the relative binding of purified IgG1 (control) versus a variety of Fc-mutant antibody technologies to immobilized human FcγRI (CD64), as determined by Surface Plasmon Resonance (SPR) assay (data courtesy of MAbsolve, https://mabsolve.com/science/#linkone). Note extremely low binding of either the YTE variant used by Paradigm (far right bar) or MAbsolve’s STR variant. Inset: The binding affinities of the non-YTE LiVE or YTE-variant LiVE-Longer ACE-2/Fusion protein chimeras to purified FcRn receptor (Ladel et al., 2018) were determined by SPR assay (Acro Biosystems). Note high affinity binding of the YTE chimera (27nM) to FcRn. See text for details.

**Figure 13.**
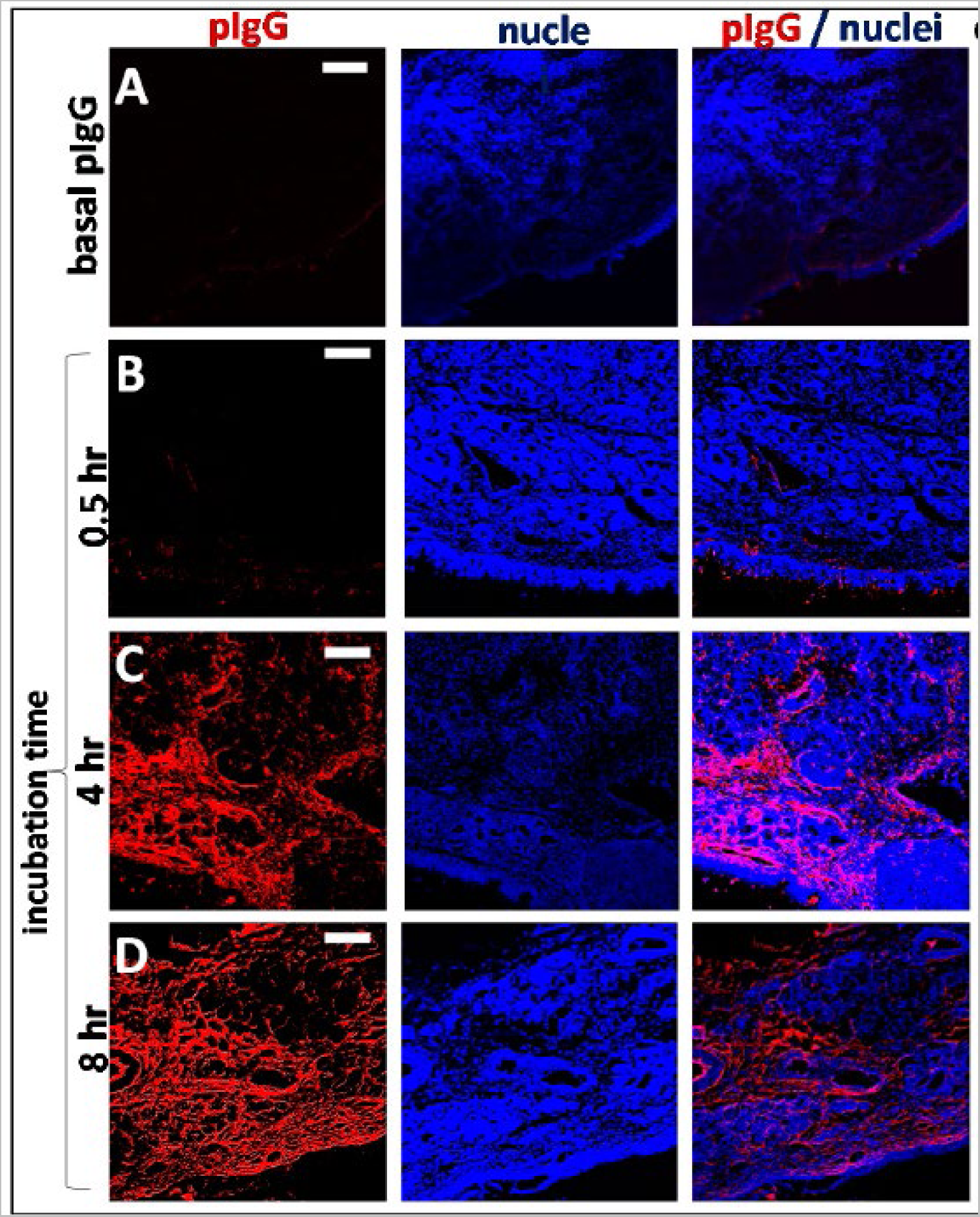
Longer half-life of nasally-applied chimeric ACE-2/Fc-silent Fusion protein is due to kinetics of FcRn-Dependent IgG Uptake into Olfactory Mucosa. Nasally-applied IgG is slowly transcytosed thru binding to FcRn receptor expressed in porcine or human olfactory epithelia. In this figure, Ladel et al. (Ladel et al., 2018) used ex vivo porcine olfactory mucosa to show to track the uptake of allogenic IgG (red) into porcine olfactory epithelia. (A) Basal levels of endogenous pIgG (porcine IgG) were detected with a low signal at the apical side, in the basal cell layer, glands, cavernous bodies, and blood vessels. This signal served as a blank and was subtracted from the photos showing the penetration of exogenous pIgG. (B) After 30 min, only the areas close to the apical side show immunoreactivity for pIgG, but some signal was detected in the lamina propria. (C) After 4 h, the pIgGs obviously distributed into the lamina propria. * indicate round structure filled with cells and mostly spared from IgG (D) After 8 h, pIgG were detected throughout the whole lamina propria. Nuclei stained with DAPI; epithelial control: quality control for tissue integrity, stained with HE. Scale bar: 100 µm. On this basis, the YTE variant of Paradigm’s ACE-2 LVE/IgG YTE chimera (which binds longer to FcRn, (see Zhu et al., 2017) is predicted to have 3-4-fold longer biological half-life when administered nasally. Reprinted with publisher’s permission from Ladel et al., Allogenic Fc Domain-Facilitated Uptake of IgG in Nasal Lamina Propria. Pharmaceutics, 10:107, 2018. See text for details.

Table I is a summary of the measured binding affinities, determined by SPR assay, of Paradigm’s LiVE and LiVE-longer chimeric constructs to purified recombinant RBD subunits, S1 subunits containing the RBD, or spike protein trimers designed to mimic the Alpha, Delta or Omicron variants of SARS-CoV-2. For comparison, the data are displayed alongside binding data for wild-type ACE-2/Fc fusion proteins from ACRO Biosystems and Genscript, which have binding affinities of 16nM and ∼3nM, respectively, to the Wuhan strain RBD. In contrast, all Paradigm constructs had low-to-mid picomolar binding affinities to the Alpha, Delta or Omicron variant protein constructs. Of special note, the highest affinity bindings were observed for the YTE variant chimera to the Omicron subvariant BA.2 spike trimer (78fM), to the Omicron subvariant BA.1 (73pM), to the Alpha B.1.1.7 variant (93pM), and for the non-YTE chimera to the Omicron variant B.1.1.529 (308nM). These data are consistent with the sVNT assays described above; implications for potential therapeutics are described in the Discussion section.

## 1.4 Discussion

This manuscript describes a new generation of chimeric ACE-2/Fusion protein hybrid molecules designed to have four important characteristics: a) ultra-high affinity binding to viral targets; b) preservation of high affinity binding across variant subgroups; c) the option of strong silencing of Fc receptor function to minimize antibody-dependent enhancement (ADE) of infection or complement antibody-dependent enhancement (C’ADE), and d) the option of binding to the FcRn receptor to increase biological half-life, particularly in upper respiratory passages. Given the rapid and consistent evolution of new variants of SARS-CoV-2 in the last two years, it is proposed that this class of chimeric molecules offers a viable new set of approaches to prophylaxis of SARS-CoV-2 and, in the future with alternate designs, other emerging viral threats yet to come.

An important feature of the design presented here is the choice of mutations to engineer into the viral receptor portion of the chimera, which for SARS-CoV-2 is the human ACE-2 protein. Multiple modeling platforms were compared and were used to choose the “L-V-E” ACE-2 variant described; at each step, the modeling results were compared favorably to recently published crystal and cryo-EM structures of ACE-2 bound to SARS-CoV-2 variants. The substitution of E90 for the wild type N eliminates the N-linked glycan, which relieves steric hindrance by the sugar and allows closer ACE-2/RBD interactions with all other mutants tested (see Figure 3). The amino acid substitutions L27 and V34, which interact with SARS-2 RBD amino acids 473 and 456 versus 455 and 453, respectively (Figure 2), were found in modeling to produce the most stabilizing ACE-2/RBD interactions (lowest D-FIRE score and K_D_ by SPR) for the widest number of SARS-2 variants, especially when paired with the E90 substitution to eliminate the N-glycosylation (See Figures 4-5 and Table I). Somewhat surprisingly, when the LVE mutant of ACE-2 was paired with the YTE sequence in the IgG portion of the chimera, the measured binding affinities to several SARS-2 variants were even greater than those measured in the non-YTE construct (to be discussed below, see Table I). Moreover, the sVNT assays reported in Figures 7-9 & 11 yielded viral neutralization data entirely consistent with the modeling and SPR binding data.

Of particular note in the context of the most recent SARS-CoV-2 Omicron sequence at the time of manuscript submission, the SARS CoV-2 RBD mutation N417 (w.t. is K417), along with other Omicron mutations viewed in 3D molecular modeling (see Figure 10, top panel), has caused this RBD amino acid to move further away from the ACE2 D30 amino acid and adopt a more vertical orientation (purple arrow next to V34), compared to the w.t. K417 which was more horizontal (not shown). Across multiple modeling simulations, the choice of valine at ACE-2 position 34 offered the widest variety of favorable ACE-2/RBD interactions, including with the recent the BA.1 or BA.2 sublineages of Omicron which have lost the K417 mutation. It is for these reasons that the chimeric molecules described here may be termed “variant agnostic”.

The intentional inclusion of the YTE variant of the antibody domain of the chimera was designed to permit increased binding of the “LiVE-Longer” chimeras to the FcRn receptor, which is known to increase the biological half-life of other IgGs currently in use by 3 to 4 fold (Borrok et al., 2017; Zhu et al., 2017). The FcRn receptor binds primarily to the CH2/CH3 interdomain area on IgG Fc, but the Fab arms also contribute to FcRn binding. For this reason, some fusion proteins such as TNFR-IgG Fc mABs (etanercept, trade name Enbrel) have a substantially shorter half-life than normal IgG (Susuki et al., 2021). For this reason, the incorporation of the YTE sequence in Paradigm’s chimeras is expected to not only saturate the FcRn widely expressed in the respiratory tract, but is predicted to substantially increase their biological half-life. In further support of this prediction, Motavizumab-YTE and Omalizumab-YTE have both been shown to have an extended half-life in healthy adults simply as a result of incorporating the YTE sequence (Robbie et al. 2013, Liu et al. 2020), a property known to be imparted by binding of this sequence to the FcRn receptor (Mackness et al., 2019). Although biological half-life has not yet been tested for the fusion proteins described here, future pharmacokinetic and pharmacodynamic studies to be performed as part of a later FDA EUA submission are expected to yield a similar half-life extension of 2-4-fold.

This is a feature that no other ACE2-Fc fusion proteins to date have taken into account, and is expected to allow lower doses, administered less frequently, to improve therapeutic efficacy. By analogy to other mAbs containing the YTE sequence (Zhu et al, 2017), it is expected that the Paradigm chimeras expressing YTE will exhibit 3-4-fold increased biological half-life, especially if administered nasally, due to high FcRn expression in the nasal and oral epithelia (Hou et al., 2020). The Surface Plasmon Resonance (SPR) data of Table I (see inset) are consistent with this hypothesis, as the YTE construct exhibited nearly 20-fold higher binding affinity to purified FcRn (27 nM) compared to the non-YTE construct (517 nM). The lower pH of the nasal cavity (∼5.5, Halwe et al., 2021) is not expected to decrease ACE-2 binding, as computational modeling of chimera-RBD binding at pH 7.4 vs 5.5 yielded DFIRE Scores of -8.54 vs. -8.01, respectively (data not shown)

In addition, and somewhat surprisingly, the LiVE-Longer YTE chimeras, when compared to their non-YTE counterparts, also showed consistently higher binding affinities to the SARS-2 protein constructs corresponding to the Alpha variant B.1.1.7 and the Omicron variants B.1.1.529 and BA.1, when these were assayed as S1 subunits or spike protein trimers (see Table I). Further, the highest affinity binding was found for the YTE chimera to the Omicron subvariant BA.2 (78 fM). Another potential benefit of incorporating the YTE sequence for FcRn binding is the presence of FcRn expressed by endothelial cells throughout the vasculature (Walters et al., 2016). Recently, extracellular vimentin expressed and released by endothelial cells was shown to act as an adjuvant to ACE-2, increasing ACE-2-mediated entry of SARS-CoV-2 into the endothelium and thereby promoting infection (Amraei et al., 2022). In light of these findings, it might be predicted that high binding of the YTE chimeras to FcRn within the vasculature, together with the increased half-life that binding imparts, would act to further inhibit vimentin-mediated ACE-2-dependent cell entry by the virus. Whether or not these hypotheses are correct will be interesting topics for future investigations.

Regardless, the new chimeric ACE-2/Fc-silent fusion proteins described here offer a promising new approach to prophylaxis of SARS-CoV-2 infection that rigorous pre-clinical testing has shown to be relatively variant-agnostic. On the basis of published data from other mAb preparations containing the YTE sequence, the biological half-life of these constructs is expected to be increased 3-4-fold above that of non-YTE fusion proteins. This feature is expected to not only increase biological half-life, but due to the high expression of FcRn in nasal and oral mucosa, enable lower and less frequent dosing of compound delivered intranasally.

Given the stability of these constructs at the acidic pH of the nasal mucosa, it is proposed that intranasal delivery or nebulization may be an optimal delivery route for this proposed prophylactic strategy against SARS-CoV-2 infection. By saturating the respiratory tract FcRn with the “LiVE Longer” mAb, the goal of these YTE (or other IgG half-life extending mutations) prophylactics is to achieve passive sterilizing immunity in future *in vivo* pre-clinical and human clinical testing. In addition, the design described here offers the possibility to exchange the ACE-2 portion of the construct with other viral receptors, in future efforts to combat viral threats that are likely to emerge.

### 1.5 Conclusions

In this report we describe the design new molecules which combine a synthetic human ACE-2 domain that is mutated to allow variant-agnostic, ultra-high affinity binding to the SARS-CoV-2 spike protein trimer. The ACE-2 domain is combined with an Fc-silent antibody domain that essentially eliminates the potential for ADI or ADE. Moreover, a third mutant option of the antibody domain of the chimera is offered, with the intent to substantially increase (3-4-fold) the biological half-life of the chimera, especially if delivered by aerosol or nasal administration. It is proposed that a nasal administration of the new chimeric molecules described herein will constitute an effective prophylactic against SARS-CoV-2 infection that will not only be effective, but also will be economically superior to current monoclonal antibody treatments for COVID-19.

### 1.6

**TABLE I.**
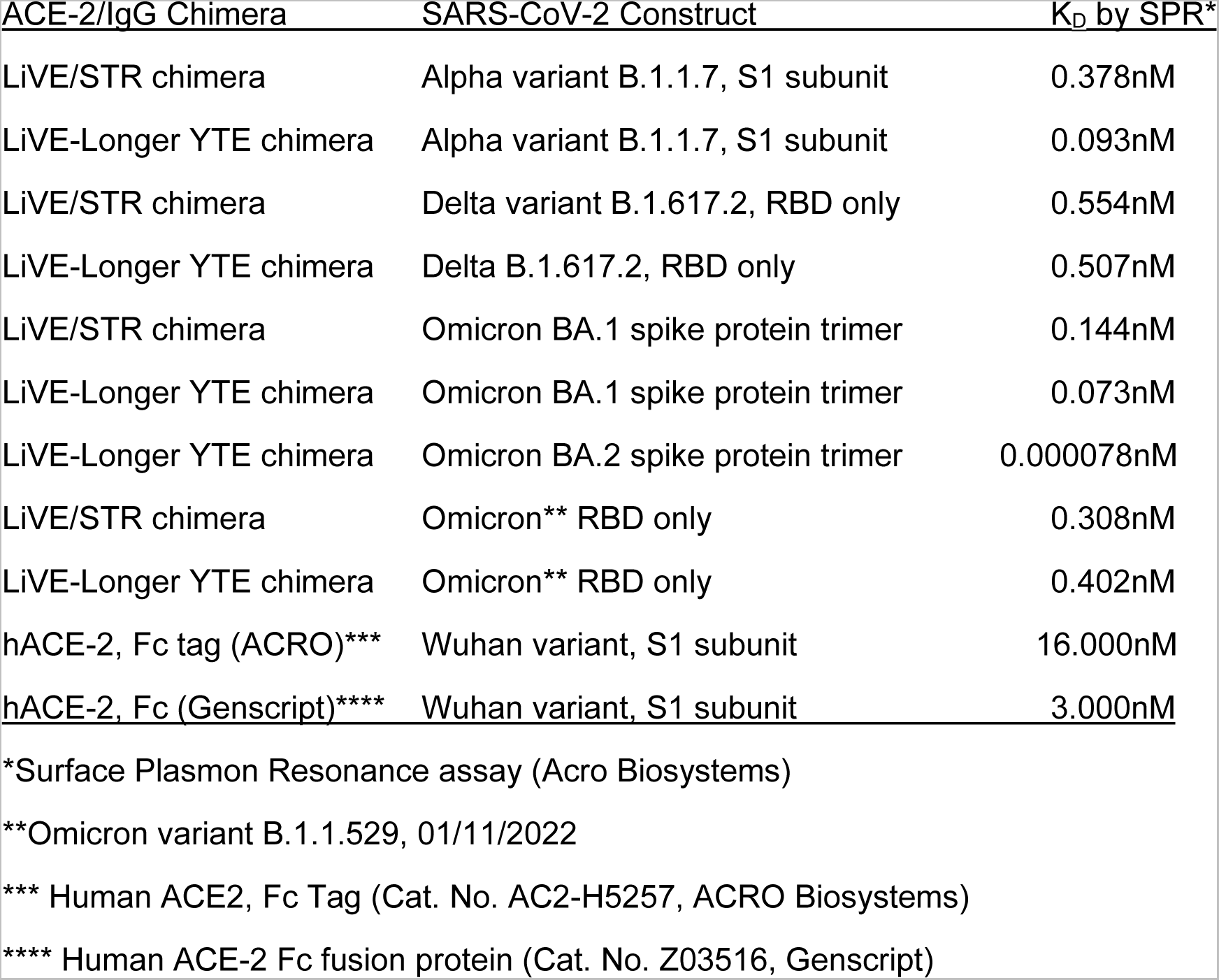
Binding Affinities of Paradigm’s ACE-2/Fusion protein Chimeras to SARS-CoV-2 Variant Constructs

| ACE-2/IgG Chimera | SARS-CoV-2 Construct | K<sub>D</sub> by SPR* |
| --- | --- | --- |
| LiVE/STR chimera | Alpha variant B.1.1.7, S1 subunit | 0.378nM |
| LiVE-Longer YTE chimera | Alpha variant B.1.1.7, S1 subunit | 0.093nM |
| LiVE/STR chimera | Delta variant B.1.617.2, RBD only | 0.554nM |
| LiVE-Longer YTE chimera | Delta B.1.617.2, RBD only | 0.507nM |
| LiVE/STR chimera | Omicron BA.1 spike protein trimer | 0.144nM |
| LiVE-Longer YTE chimera | Omicron BA.1 spike protein trimer | 0.073nM |
| LiVE-Longer YTE chimera | Omicron BA.2 spike protein trimer | 0.000078nM |
| LiVE/STR chimera | Omicron** RBD only | 0.308nM |
| LiVE-Longer YTE chimera | Omicron** RBD only | 0.402nM |
| hACE-2, Fc tag (ACRO)*** | Wuhan variant, S1 subunit | 16.000nM |
| hACE-2, Fc (Genscript)**** | Wuhan variant, S1 subunit | 3.000nM |

\*Surface Plasmon Resonance assay (Acro Biosystems)
\*\*Omicron variant B.1.1.529, 01/11/2022
\*\*\* Human ACE2, Fc Tag (Cat. No. AC2-H5257, ACRO Biosystems)
\*\*\*\* Human ACE-2 Fc fusion protein (Cat. No. Z03516, Genscript)

